# Quercetin handles cellular oxidant / antioxidant systems and mitigates immunosenescence hallmarks in human PBMCs: an *in vitro* study

**DOI:** 10.1101/2021.10.23.465570

**Authors:** Samia Bouamama, Amina Bouamama

## Abstract

Immunosenescence, oxidative stress, and low vaccine efficacy are important symptoms of aging. The goal of our study was to see if quercetin had anti-aging and stimulating effects on PBMC immune cells in vitro. In the presence of concanavalin a, PBMCs were isolated from healthy elderly and young people and cultured in a complete RPMI-1640 medium supplemented with quercetin. Cell proliferation was assessed using the MTT colorimetric assay after a 48-hour incubation period. Spectrophophotometric assays were used to assess oxidative biomarkers (PC, MDA, and GSH). The ELISA method was used to determine the amount of IL-2 released. A Griess reagent was used to investigate *i*NOS activity. When compared to young control cells, aged PBMCs had lower proliferation potency, lower IL-2 and NO release, and higher MDA and PC levels. Importantly, quercetin-treated aged PBMCs have a high proliferative response comparable to young cells, restored *i*NOS activity, and increased levels of GSH antioxidant defences. In comparison to untreated aged PBMCs, treated PBMCs have lower lipo-oxidative damage but higher PC levels. Quercetin may be used as a promising dietary vaccinal adjuvant in the elderly, it has significant effects in reducing immunosenescence hallmarks, as well as mitigating the lipo-oxidative stress in PBMCs cells.

**Practical abstract:** Quercetin is a common dietary polyphenol found in fruits and vegetables that is well known for its antioxidant activity in radical scavenging and anticancer properties characterized by immune system stimulation. *In vitro*, quercetin boosts immune responses in the elderly and reduces immunosenescence symptoms, according to our findings. As a result, this molecule could be a safe and promising vaccine adjuvant for boosting immunity and reducing aging complications in immune cells as well as reducing oxidative stress in PBMC cells.

## 1. Introduction

Aging is a complex and multietiological process that involves gradual and spontaneous biochemical and physiological changes, such as increased susceptibility to diseases, exposure to harmful environmental conditions, and a loss of mobility and agility [1]. The body’s major functions, such as the circulatory and digestive systems, the endocrine system, and the immune system, deteriorate over time. Similarly, one of the signs of aging is the deterioration of cognitive and memory functions [2].As people get older, the quantitative and functional characteristics of various types of immune cells deteriorate, resulting in a proclivity for inflammation, increased susceptibility to infectious diseases, a decreased ability to maintain tolerance to self-antigens, and a reduced response to immunisation [3,4].Previous research has found a strong link between immune function and age, with older people’s PBMC cell proliferation being lower than that of younger people [5]. As a result, when compared to other age groups, older adults are more susceptible to infectious diseases that are more frequent and last longer. As a result, according to the US Centers for Disease Control and Prevention (CDC), older adults in the United States accounted for more than 92.45% of COVID-19 deaths in the country as of December 2020 [4]. Despite the significant progress made with current vaccines, the majority of vaccines still fail to provide long-term immunity in older adults [6]. Natural adjuvants, for example, have been accepted as effective strategies for increasing the immunogenicity of existing vaccines proposed in recent years, and they may provide effective protection for older adults [4]. Plant-derived adjuvants are relatively non-toxic and do not cause significant side effects, which is a major concern with synthetic compounds; second, they have been shown to potentiate the immune response, making their use in the development of vaccines appealing [7]. Natural dietary polyphénols, particularly quercetin, have been shown to have profound effects on immunity, according to research literature, and thus could be used as adjuvants [8]. Quercetin, 3,3’,4’,5,7-pentahydroxyflavone, is a widely distributed secondary metabolite in plants that is one of the most abundant vegetal flavonoids in the human diet. Apples, onions, berries, leafy green vegetables, hot peppers, red grapes, and black tea are all good sources of antioxidants. Quercetin has antioxidant, free radical–scavenging, and anti-inflammatory properties that could affect immune system competence and pathogen resistance [9]. Existing research shows that quercetin has senescence-delaying activity in primary cells and rejuvenating effects in senescent cells in vitro [10]. Quercetin exhibits adjuvant activity by enhancing Th2 immune response in ovalbumin-immunised mice, according to previous studies demonstrating the results of immunisation with plant phenolics [8]. In the present study, we assessed in vitro quercetin treatment-induced changes on PBMCs cell proliferation, pro-/anti-oxidant state, IL-2, and nitric oxide release from both young and elderly subjects to better understand the possible clinical uses and safety of quercetin as vaccinal adjuvants, particularly for the elderly.

## 2. Methods

### 2. 1. Reagents

Con a : Concanavalin A, MTT: (3-(4,5-dimethyl-2-thiazolyl)-2,5-diphenyl-2H-tetrazolium bromide), Histopaque 1077, DNPH (2,4-dinitrophenyl hydrazine), DTNB: (5,5-dithiobis-2- nitrobenzoic acid), Trypan blue dye and all other chemicals were purchased from Sigma-Aldrich company (Sigma, St Louis, MO, USA). Incomplete Roswell Park Memorial Institute 1640 (RPMI 1640) medium was produced by Gibco Life Technologies Inc. (Paisley, U.K). it was aseptically supplemented with 25 mM HEPES buffer, 10% heat-inactivated fetal calf serum, L-glutamine (2 mM), 2-mercaptoethanol (5 × 10^−5^ M), penicillin (100 UI/ml) and streptomycin (100 μg/ml).

Quercetin (Sigma Aldrich, France) was dissolved in DMSO (<1% in culture medium) to make a stock solution (200 μg/ml) ; the final concentration of quercetin in the culture medium was 20 μg/ml. DMSO concentrations less than 1% were previously found to be neither cytotoxic nor genotoxic to mammalian cells.

### 2.2. Study population

PBMCs cells were isolated from peripheral blood of 16 healthy donors (10 elderly women and men and 06 young healthy subjects ; In brief, fasting venous blood were collected from healthy volunteers (young subjects aged between 20 to 33 years old) and the elderly (aged over than 65 years old). All patients in this study are informed about the purpose of the work according to the statement of Helsinki 1964 and no investigation is conducted without prior consent signed by the participants. This is achieved by respecting the anonymity and confidentiality of information. The study was performed under the approval of the Ethics Committee of Tlemcen University Hospital.

### 2.3. PBMCs culture *in vitro*

The gradient density centrifugation method was used to obtain blood PBMCs from healthy donors [11], In brief; 4 ml of whole EDTA blood was layered into 3 ml of Histopaque 1077 solution and centrifuged at 4000 rpm for 40 minutes. The cells at the plasma-Histopaque interface were collected and washed twice with balanced saline solution (HBSS), and the cell clot was kept in RPMI complete medium. The Trypan blue exclusion method was used to determine cell viability. In RPMI-1640 medium, 4.10^6^ PBMCs were seeded aseptically in a 96-well cell culture plate in the presence of 5 μg/ml of the mitogen Con A and 20 μg/ml of quercetin. Negative controls included only solvent treatment. At 37°C, the cell culture was incubated in a humidified incubator with 5% CO2.

### 2.4. PBMCs cellular proliferation assay

Cell proliferation was analyzed with cell viability assays that measure the rate of cellular metabolism with MTT substrate as described earlier [12]. Cell viability was assessed with an additional 10 μl/well of MTT (5 mg/ml) after 48 hours of incubation. At a wavelength of 570 nm, the plates were read on a microplate reader. Proliferation of PBMCs is measured by the ratio of treated cells to untreated cells that served as controls, expressed as a percentage of absorbance. Alternatively:

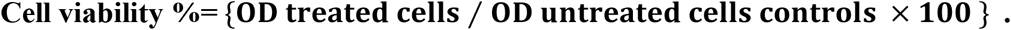

### 2.5. Determination of intracellular redox biomarkers

The concentration of MDA in cell homogenate was measured using the thiobarbituric acid reactive species assay described by **Nourooz –Zadeh et al. (1996)** to determine the level of lipid peroxidation in PBMCs cells [13]. The 2,4-dinitrophenyl hydrazine (DNPH) reaction was used to measure carbonyl proteins (markers of protein oxidation) in cellular homogenate. [14]. as previously described, the reduced glutathione GSH level in PBMCs was measured using 5,5-dithiobis-2-nitrobenzoic acid (DTNB) [15].

### 2.6. IL-2 cytokine Elisa –assay

An enzyme-linked immunosorbent assay (ELISA) kit was used to determine the levels of IL-2 cytokine released in PBMC free supernatants (Abfrontier, Multiplex Human Cytokine ELISA Kit).

### 2.7. Nitric oxide release in cells supernatants (*i*NOS activity)

The method described by Guevara et al. **(1998)** was used to check NO release by PBMCs [16]. This is based on the concentration of whole nitrites in the medium. The obtained supernatant is mixed with the Griess reagent (sulfanilamide and N-1naphtylethylenediamine dihydrochloride in orthophosphoric acid) after centrifugation. The intensity of the pink-rose coloration is proportional to the NO concentration. Optical densities are measured at 540 nm using a spectrophotometer. NO concentrations are plotted on a calibration curve made with sodium nitrite NaNO2 (0-100M).

### 2.8. Statistics

Minitab 16 statistical software and Microsoft Excel 2007 were used to conduct all statistical analyses. The data is presented as a mean± SD. The one-way ANOVA test was used to determine differences between groups, followed by the Turkey grouping method. At least three times, all in vitro cultures were repeated. The threshold for statistical significance was set at *p* < 0.05.

## 3. Results

### 3.1. In vitro effect of quercetin on cell proliferation of PBMCs

When compared to young controls, the proliferative potency of PBMCs from the elderly is significantly lower (Fig1). In vitro, quercetin treatments restored cellular proliferation in the elderly in the same way that they did in young subjects (p=0,029).

**Fig 1:**
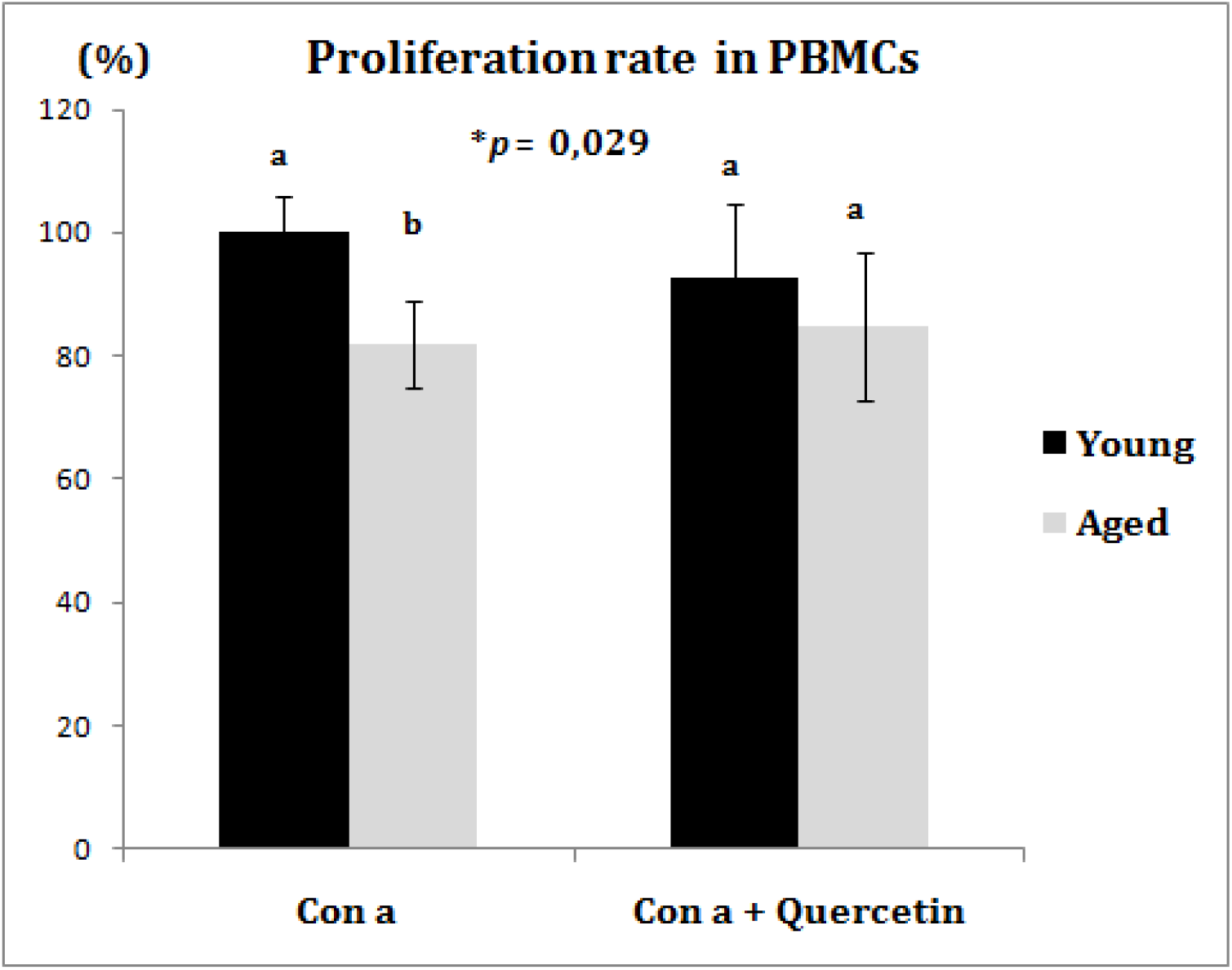
PBMCs cellular proliferation rate in response to quercetin and Con a treatments. Each value represents the mean ± SD. Significant differences between groups are indicated by small letters (a,b,c,d), with *P-* One-way ANOVA less than 0.05. PBMCs: peripheral Blood Mononuclear Cells, Con A: concanavalin a lectin.

### 3.2. Intracellular oxidant/antioxidant biomarkers in PBMCs

MDA and PC levels in the PBMCs of elderly subjects were significantly higher than in young controls. Quercetin treatment resulted in a significant decrease in lipid peroxides in both young and old PBMCs (Fig 2), but with an increases in PC only in aged PBMCs (**p = 0,0008) (Fig 3) When PBMCs from older subjects were compared to PBMCs from young controls, they showed similar GSH cellular levels, whereas quercetin treatment significantly increased intracellular GSH amounts only in the older subjects (Fig 4).

**Fig 2:**
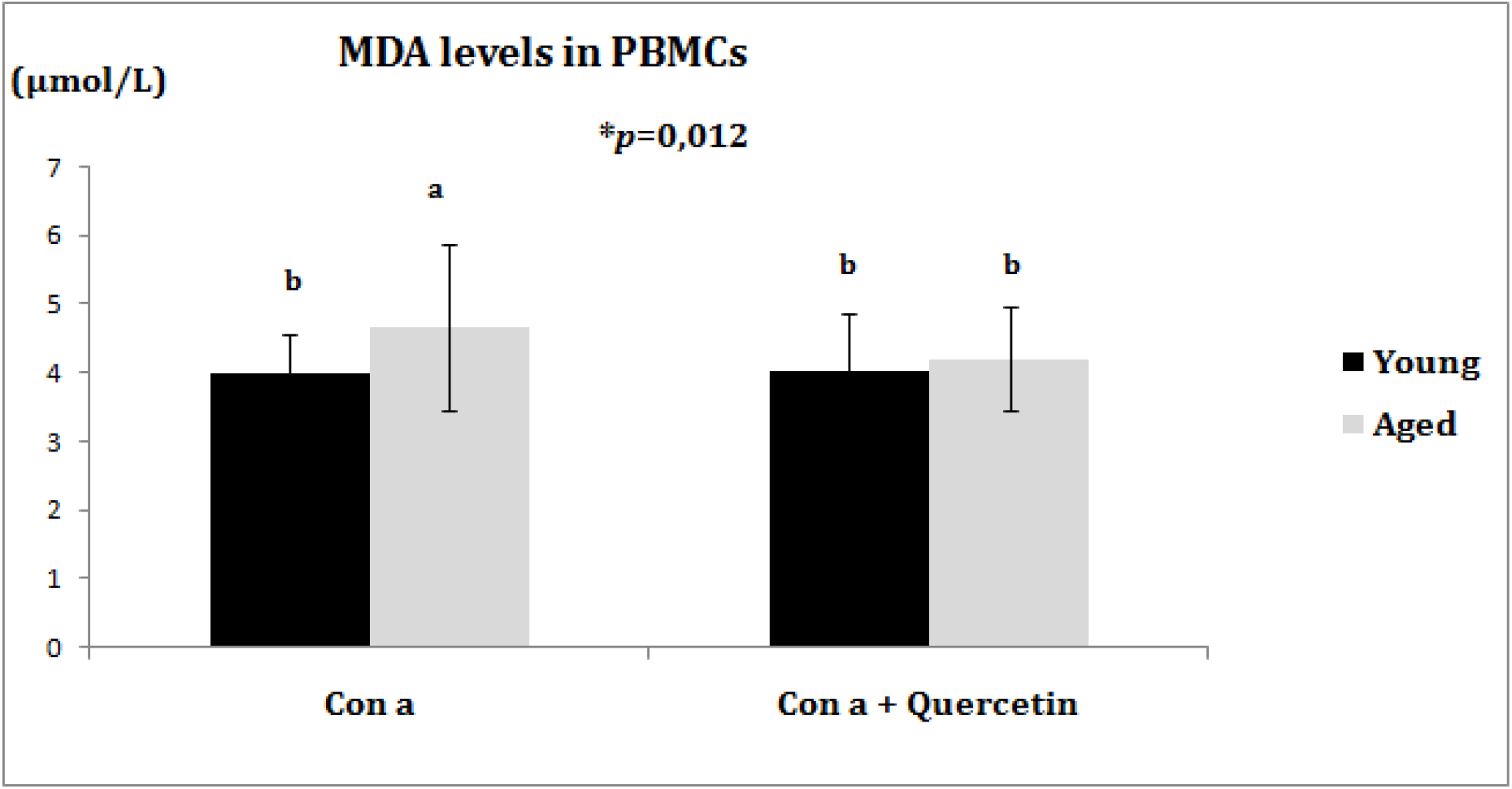
Intracellular Malondialdehydes levels in PBMCs. Each value represents the mean ± SD. Significant differences between groups are indicated by small letters (a,b,c,d), with *P-* One-way ANOVA less than 0.05. PBMCs: peripheral Blood Mononuclear Cells, Con A: concanavalin a lectin. MDA: Malondialdehydes.

**Fig 3:**
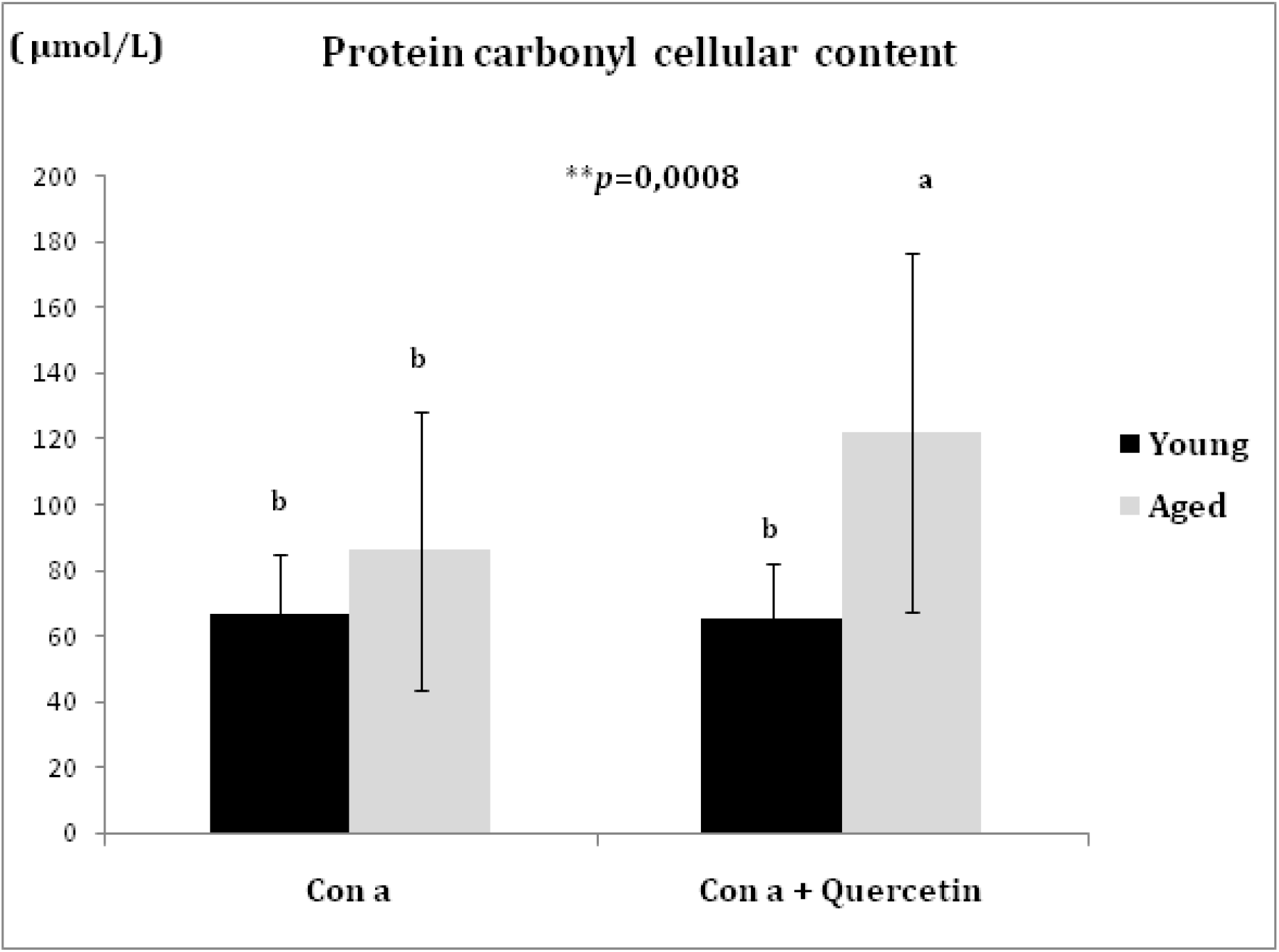
Intracellular protein carbonyl content in PBMCs. Each value represents the mean ± SD. Significant differences between groups are indicated by small letters (a,b,c,d), with *P-* One-way ANOVA less than 0.05. PBMCs: peripheral Blood Mononuclear Cells, Con A: concanavalin a lectin.

**Fig 4:**
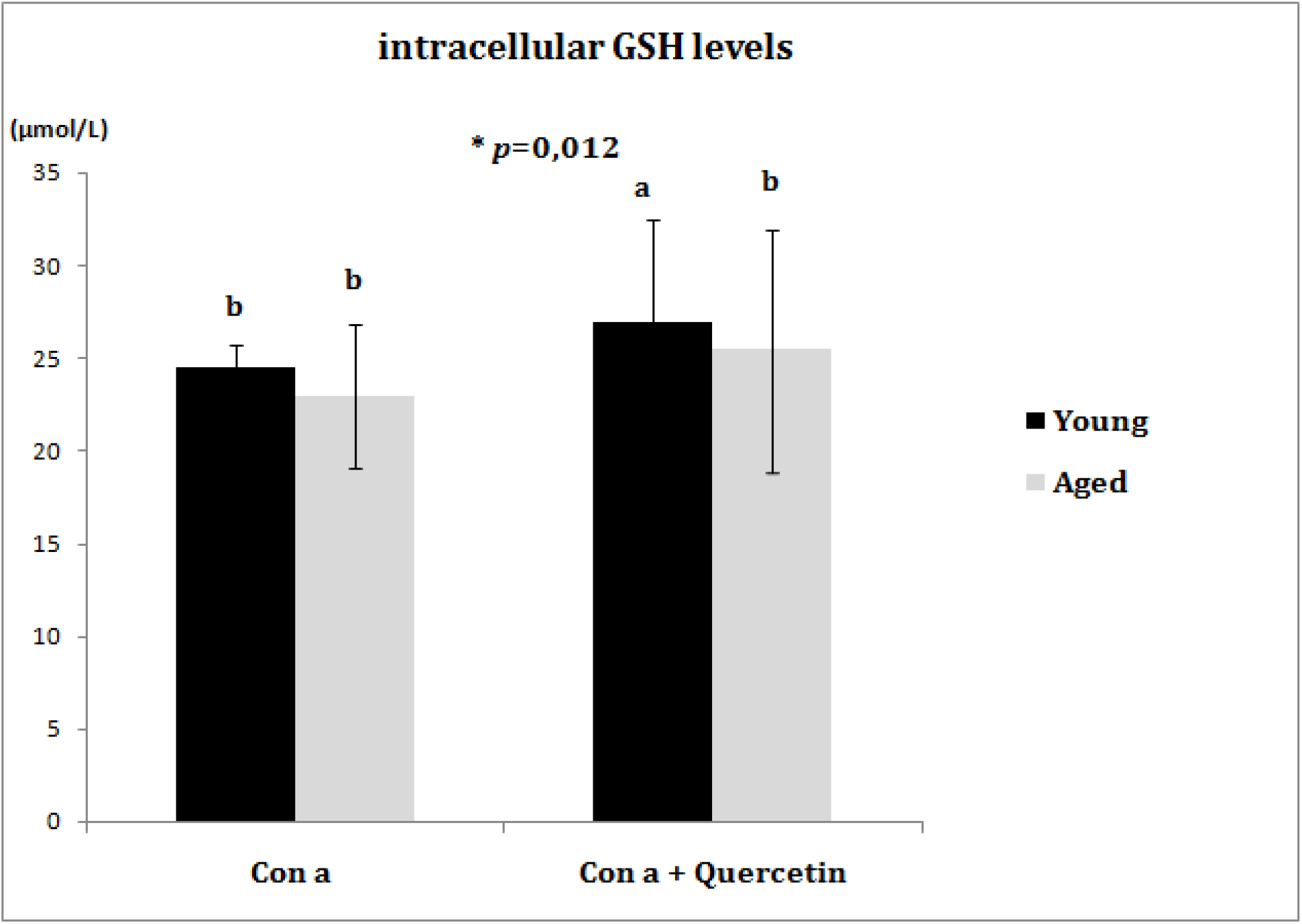
Intracellular GSH contents in PBMCs. Each value represents the mean ± SD. Significant differences between groups are indicated by small letters (a,b,c,d), with *P-* One-way ANOVA less than 0.05. PBMCs: Peripheral Blood Mononuclear Cells, Con A: concanavalin a lectin.

### 3.3. IL-2 and *i*NOS activity in PBMCs

When compared to young controls, the elderly subjects’ IL-2 cellular release was significantly reduced. Quercetin treatment reduces IL-2 levels in both young and aged cells (**p= 0, 0006). The aged subjects’ PBMCs released less NO cellular content, implying lower *i*NOS activity, than the control subjects’ PBMCs. Quercetin treatment of PBMCs cells significantly increased *i*NOS activity in the aged cells (Table 1).

**Table 1.**
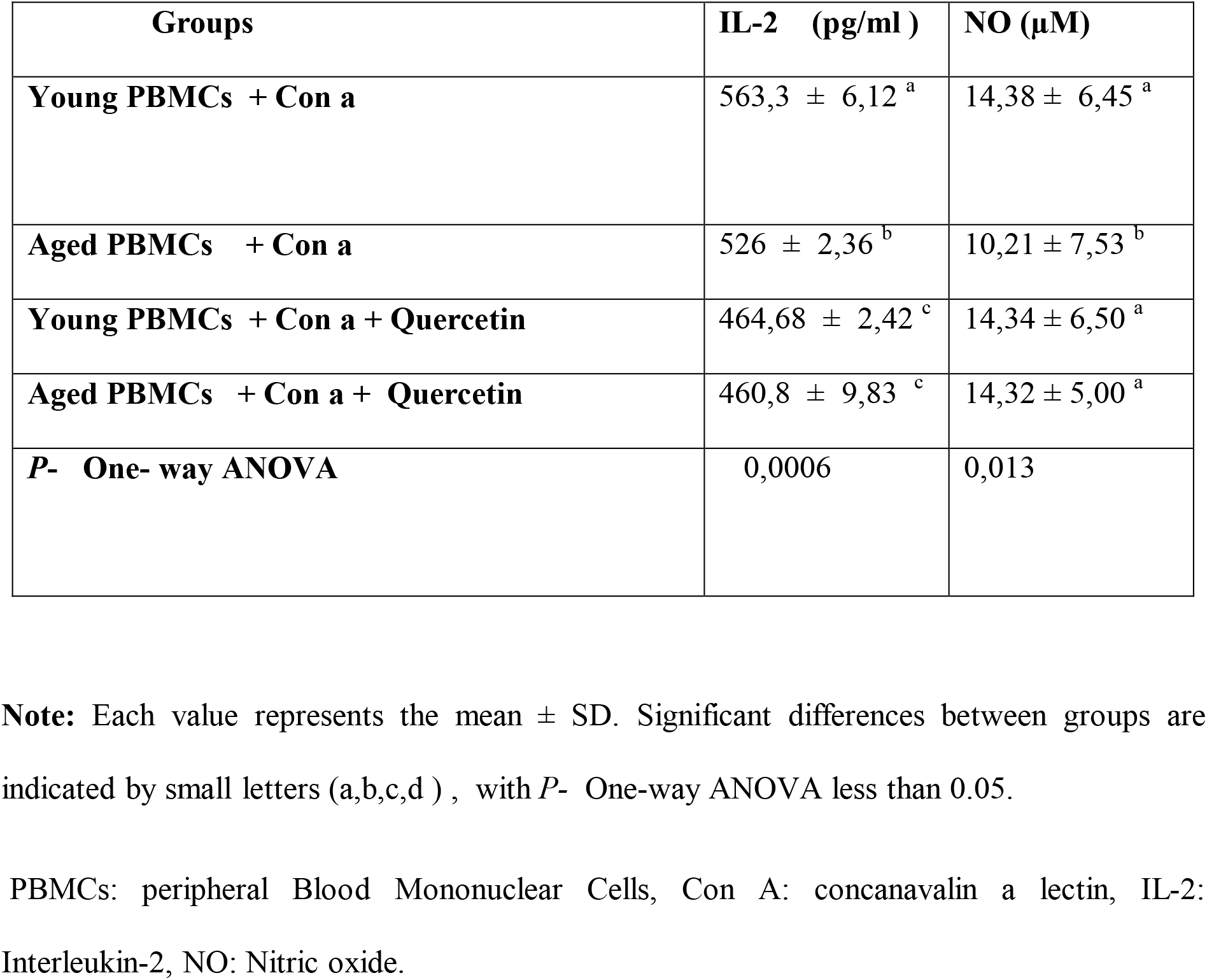
IL-2 and NO releases from PBMCs.

| **Groups** | **IL-2 (pg/ml )** | **NO (μM)** |
| --- | --- | --- |
| **Young PBMCs + Con a** | 563,3 ± 6,12 <sup>a</sup> | 14,38 ± 6,45 <sup>a</sup> |
| **Aged PBMCs + Con a** | 526 ± 2,36 <sup>b</sup> | 10,21 ± 7,53 <sup>b</sup> |
| **Young PBMCs + Con a + Quercetin** | 464,68 ± 2,42 <sup>c</sup> | 14,34 ± 6,50 <sup>a</sup> |
| **Aged PBMCs + Con a + Quercetin** | 460,8 ± 9,83 <sup>c</sup> | 14,32 ± 5,00 <sup>a</sup> |
| ***P*- One- way ANOVA** | 0,0006 | 0,013 |

**Note:** Each value represents the mean ± SD. Significant differences between groups are indicated by small letters (a,b,c,d) , with *P*- One-way ANOVA less than 0.05.
PBMCs: peripheral Blood Mononuclear Cells, Con A: concanavalin a lectin, IL-2: Interleukin-2, NO: Nitric oxide.

## 4. Discussion

The primary physiological role of the human immune system is to eliminate potentially harmful compounds and a wide range of potentially threatening organisms; it has evolved to include many different cell types, many communicating molecules, and multiple functional responses. It is self-evident that a healthy immune system is required for effective defence against pathogenic organisms. thus, people with weakened immune systems are more likely to become infected and have more serious infections [17]. Unfortunately, as people get older, they develop immunosenescence, which is characterised by a reduced response to antigens and vaccines [6]. The current study backs up our previous findings, which show that when PBMC is stimulated solely by Con a, proliferation is significantly reduced in elderly people compared to young controls. The immunosuppressive mechanisms involve a series of cellular and molecular events involving the alteration of several biochemical pathways and cell populations [3]. Importantly, Con a in combination with quercetin significantly increased PBMC proliferation in both elderly and young cells. In PBMCs, quercetin appears to stimulate cellular proliferation by acting on the oestrogen receptor (ER), and evidence suggests that physiologically relevant quercetin concentrations can exert phytoestrogen-like activity [18]; it can also activate GC-A or other isoforms of guanylate cyclase. [19]. Several age-related changes in immune functions and impaired cell proliferation may be linked to oxidative stress, according to Harman’s free radicals theory of ageing. At the cellular level, excessive amounts of ROS cause damage to cellular components such as carbohydrates, polyunsaturated fatty acids, DNA, and proteins; thus, excessive amounts of ROS cause damage. [20]. It is well known that the concentration of oxidised biomolecules, such as MDA and PC, rises with age. [21]. When a hydrogen atom is removed from the methyl group of an unsaturated fatty acid, oxidised lipids form. MDA is a keto-aldehyde formed when unsaturated lipids are peroxidized. Protein peroxidation occurs when excess MDA reacts with free amino acids. The findings of our study are consistent with **Dalle-Donne et al**., **(2006)** [22]; We observed an increase in two pro-oxidant parameters in aged PBMCs cells: MDA and PC, but no decrease in cellular antioxidant potency GSH when compared to young PBMCs. Increased cellular oxidative stress plays an important role in immunosenescence and cellular aging [23].Changes in the permeability of the cell membrane may occur when lipid peroxidation is increased.

In fact, PBMCs from elderly patients were exposed to intracellular oxidative stress and had higher levels of oxidant biomarkers in our study. Several in vivo and in vitro studies have found that dietary antioxidant molecules can reduce tissue damage caused by oxidative stress [24]. Our findings showed that in the presence of quercetin, MDA levels in aged PBMCs decreased significantly; on the other hand, PC levels increased dramatically in aged PBMCs treated with quercetin. Because polyphenols are ambivalent compounds, previous published reports showed that quercetin acts as a double-edged sword in the cell. The same chemical properties that make them anti-oxidants can also make them prooxidants under certain circumstances [25]. As a result, quercetin, like the majority of polyphénols, can be oxidised to o-quinone derivatives that can catalyse oxidative deamination of primary amines. Indeed, pyrroloquinoline quinone, an o-quinone derivative, is known to be an efficient mediator of the oxidation of primary amines in proteins, such as the ε-amino group of lysine residues, to form α-aminoadipic-5-semialdehyde oxidative products with carbonyl functionality [26].Because it acquires potent prooxidant properties in the presence of transition metals (iron and/or copper), which are frequently accumulated in aged and senescent cells, we noticed that quercetin acts on PBMCs in an age-dependent manner. It has potent prooxidant effects in aged cells more than young cells [27, 28].

Quercetin is best known for its antioxidant properties, despite the fact that it has been shown to have a prooxidant effect under certain physiological conditions. Our findings support previous research that suggests quercetin boosts cellular antioxidant defences by increasing GSH levels in cells. GSH is an important cellular defence mechanism against oxidative injury, and its depletion causes mitochondrial damage as a major side effect.[29].

In this study, we found that after quercetin treatment, young PBMCs have higher GSH intracellular levels than untreated cells, in contrast to aged cells. Low GSH levels in aged cells compared to young cells strongly suggest that ROS/RNS have depleted GSH in aged cells and that there is a high cellular demand for GSH to cope with remarked oxidative stress.

T lymphocytes, a subset of PBMCs, play a critical role in the immune response to specific antigens. They secrete a panoply of cytokines among them IL-2, its primary effect is to induce T-cell proliferation and differentiation. IL-2 increases cytokine synthesis, potentiates Fas-mediated apoptosis, and promotes regulatory T cell development. It causes also proliferation and activation of NK cells and B-cell and antibody synthesis [30].

IL-2 is a crucial interleukin that promotes T lymphocyte clonal expansion. IL-2 release has been shown to decrease with age in previous studies. [11, 31]. Quercetin downregulates the release of IL-2 in both young and old people. Our findings are consistent with previous research, indicating that quercetin inhibits IL-2 cytokine production via an IL-2R alpha-dependent mechanism. [32].

Nitric oxide (NO) is another signalling molecule that is involved in skeletal muscle contraction, endothelial function, mitochondrial biogenesis and respiration, muscle repair, and antioxidant defences [33]. Among other physiological processes, antimicrobial cytotoxicity has also been linked to it for a variety of microorganisms, including bacterial, parasitic, fungal, and viral pathogens, as well as tumor cells. [34]. the use of NO as an immunity booster is supported by the literature, so nitric oxide-releasing compounds could be beneficial in this regard. [35]. Quercetin in combination with Con a increased NO release from PBMCs in both elderly and young subjects. Polyphenols may enhance NO production via the eNOS expression pathway and activity, according to previous literature data; these molecules also promote NO bioavailability via their antioxidant effects, which protect NO from breakdown by ROS. [33].

Quercetin has potent immunostimulatory properties and has been shown to reduce the effects of ageing in immune cells. While also reducing oxidative stress in PBMCs.

## 5. Conclusion and study limits

According to our promising preliminary findings, quercetin appears to have potential as a dietary vaccine adjuvant and may be used safely to boost immunity in the elderly. however, we cannot easily determine which vaccines quercetin is suitable for and/or whether quercetin can be used as a universal vaccine adjuvant. As a limitation, it is important to note that, as with any vaccination strategy, the efficacy of an adjuvant molecule should be studied in specific antigen-adjuvant combinations, necessitating future clinical trials.

## Abbreviations

Con a: Concanavalin a
GSH: Reduced glutathione
IL-2: Interleukin-2
iNOS: Inducible Nitrite oxide synthase
MDA: Malondialdehydes
NO: Nitric oxide
OD: Optical density
PBMCs: Peripheral blood mononuclear cells
PC: proteins carbonyl

## Acknowledgements

We are warmly thankful to the Algerian Ministry of Higher Education and Scientific Research and DGRSDT for their financial support, and we thank the volunteers and all the personnel of the Ppabionut research laboratory for their scientific collaboration. The authors are grateful to Pr. Hafida Merzouk and Dr. Farid Berroukech for supplying the quercetin molecule and other chemical reactants.

## Authors’ contributions

Dr. Samia Bouamama designed the study, contributed to running the laboratory work, analysed the data, and drafted the paper. Amina Bouamama contributed to the critical reading and English language editing of the manuscript. All the authors have read the final manuscript and approved the submission.

